# The speed of human social interaction perception

**DOI:** 10.1101/579375

**Authors:** Leyla Isik, Anna Mynick, Dimitrios Pantazis, Nancy Kanwisher

## Abstract

The ability to detect and understand other people’s social interactions is a fundamental part of the human visual experience that develops early in infancy and is shared with other primates. However, the neural computations underlying this ability remain largely unknown. Is the detection of social interactions a rapid perceptual process, or a slower post-perceptual inference? Here we used magnetoencephalography (MEG) decoding and computational modeling to ask whether social interactions can be detected via fast, feedforward processing. Subjects in the MEG viewed snapshots of visually matched real-world scenes containing a pair of people who were either engaged in a social interaction or acting independently. The presence versus absence of a social interaction could be read out from subjects’ MEG data spontaneously, even while subjects performed an orthogonal task. This readout generalized across different scenes, revealing abstract representations of social interactions in the human brain. These representations, however, did not come online until quite late, at 300 ms after image onset, well after the time period of feedforward visual processes. In a second experiment, we found that social interaction readout occurred at this same latency even when subjects performed an explicit task detecting social interactions. Consistent with these latency results, a standard feedforward deep neural network did not contain an abstract representation of social interactions at any model layer. We further showed that MEG responses distinguished between different types of social interactions (mutual gaze vs joint attention) even later, around 500 ms after image onset. Taken together, these results suggest that the human brain spontaneously extracts the presence and type of others’ social interactions, but does so slowly, likely relying on iterative top-down computations.

## Introduction

As fundamentally social primates, humans need to know who is doing what to whom, and why. Indeed, the ability to perceive and interpret social interactions between other agents develops early in infancy (Hamlin et al., 2007), is shared with other primates (Sliwa and Freiwald, 2017), and is apparently computed in a specialized region of the posterior superior temporal sulcus (Isik et al., 2017; Walbrin et al., 2017). These findings underscore the importance of social interaction perception, but leave unanswered the question of how this information is extracted from visual input. In particular, is social interaction recognition a rapid feedforward process, akin to object recognition, or a slower post-perceptual inference?

Considerable evidence suggests that much of visual perception in primates, including face, scene, and “core” object recognition, is computed by rapid and largely feedforward pattern classification processes. First, these tasks in primates are well approximated by purely feedforward neural network models, not only in terms of accuracy but also in terms of the representations extracted (Khaligh-Razavi and Kriegeskorte, 2014; Radoslaw Martin Cichy, 2016; Yamins et al., 2014). Second, visual recognition in primates is fast, occurring within 200ms of image onset, as expected of a largely feedforward process. These fast latencies have been demonstrated for face (Bentin et al., 1996; Dobs et al., 2018), scene (Cichy et al., 2016a; Greene and Hansen, 2018), and object (Carlson et al., 2013a; Isik et al., 2014; Yamins et al., 2014) recognition. In contrast, some visual information cannot be computed from bottom-up visual information alone. Object recognition under complex viewing conditions, such as occlusion takes longer (~300 ms), and cannot be performed with purely feedforward models (Rajaei et al., 2018; Tang et al., 2018, 2014). Generative models offer an attractive solution to these challenging vision problems (Wu et al., 2016; Yuille and Kersten, 2006). Rather than relying solely on bottom-up cues, these systems build models of objects and the world around them, and use these generated models as hypotheses to interpret incoming visual information.

Behavioral studies have suggested that the perception of social interactions shares some of the hallmarks of a classic visual pattern recognition problem, face recognition. First, people are better able to perceive social interactions when stimuli are presented upright rather than inverted, but the same is not true for perception of independent actions (Papeo et al., 2017). Second, social interactions receive preferential access to visual awareness (Su et al., 2016) and facilitate visual processing of groups of individuals (Vestner et al., 2019). Others have tried to address this question with computational modeling. One study showed that different types of social interactions can be distinguished based on bottom-up visual cues (Blythe et al., 1999), but more recent work has suggested that top down or generative models are required to solve this problem (Ben-Yosef et al., 2017; Ullman et al., 2009). Importantly, all these modeling efforts focused on *categorizing* different types of social interactions, and it remains an open question whether feedforward computations are sufficient to *detect* social interactions.

Here we used MEG decoding and computational modeling to ask whether social interactions can be detected, in either humans or machines, via fast, feedforward processing. Using decoding methods, we ask whether the detection of a social interaction in a visual stimulus occurs on the rapid time scale of invariant object recognition (about 150 ms), as predicted from a feedforward pattern classification model, or more slowly, as expected if it requires a top-down inference. Here we find both the presence and type of social interaction could be decoded from subjects’ MEG data, but this readout occurred quite late, at 300 ms and 500 ms for detection and categorization, respectively. In a second experiment, we showed that this readout did not occur earlier even when subjects performed an explicit social interaction detection task. Finally, using deep convolutional neural networks we tested whether a standard feedforward computational model could distinguish between scenes with vs. without a social interaction. Consistent with our neural timing data, we found that a purely feedforward model could not distinguish between scenes with vs. without a social interaction.

## Results

The below analyses were pre-registered on the Open Science Framework platform: https://osf.io/3vnem/registrations. Any deviations from our pre-registration are noted as exploratory analyses.

### Experiment 1

#### Late, spontaneous readout of social interactions

To identify MEG signals that contain information about the presence of social interactions, sixteen naïve subjects viewed visually matched images of different actor pairs in one of five different social or non-social conditions shot in one of 12 different scenes (60 total images, **Figure 1A-C**). The five conditions were: 1) joint attention (two actors looking at the same object, a classic form of social interaction), 2) mutual gaze (two actors looking at each other, a different form of social interaction), 3) independent action 1 (two actors engaged in separate independent actions, i.e., no social interaction), 4) independent action 2 (a different instance of the two engaged in separate independent actions, no social interaction), 5) watch (one actor watching the other who is looking away, a one-way interaction or “perceptual access”). We defined social interactions broadly to include either joint attention or mutual gaze. In a post-MEG behavioral experiment, subjects rated the joint attention and mutual gaze images as significantly more social than the independent action images (**Figure 1D**, p =1.9×10^−16^, two-sided t-test) and also slightly more interesting (**Figure 1E**, p = 0.003). Mutual gaze images were rated as slightly more social than joint attention images (p = 0.017), but there was no difference in their interest rating (p = 0.41).

**Figure 1:**
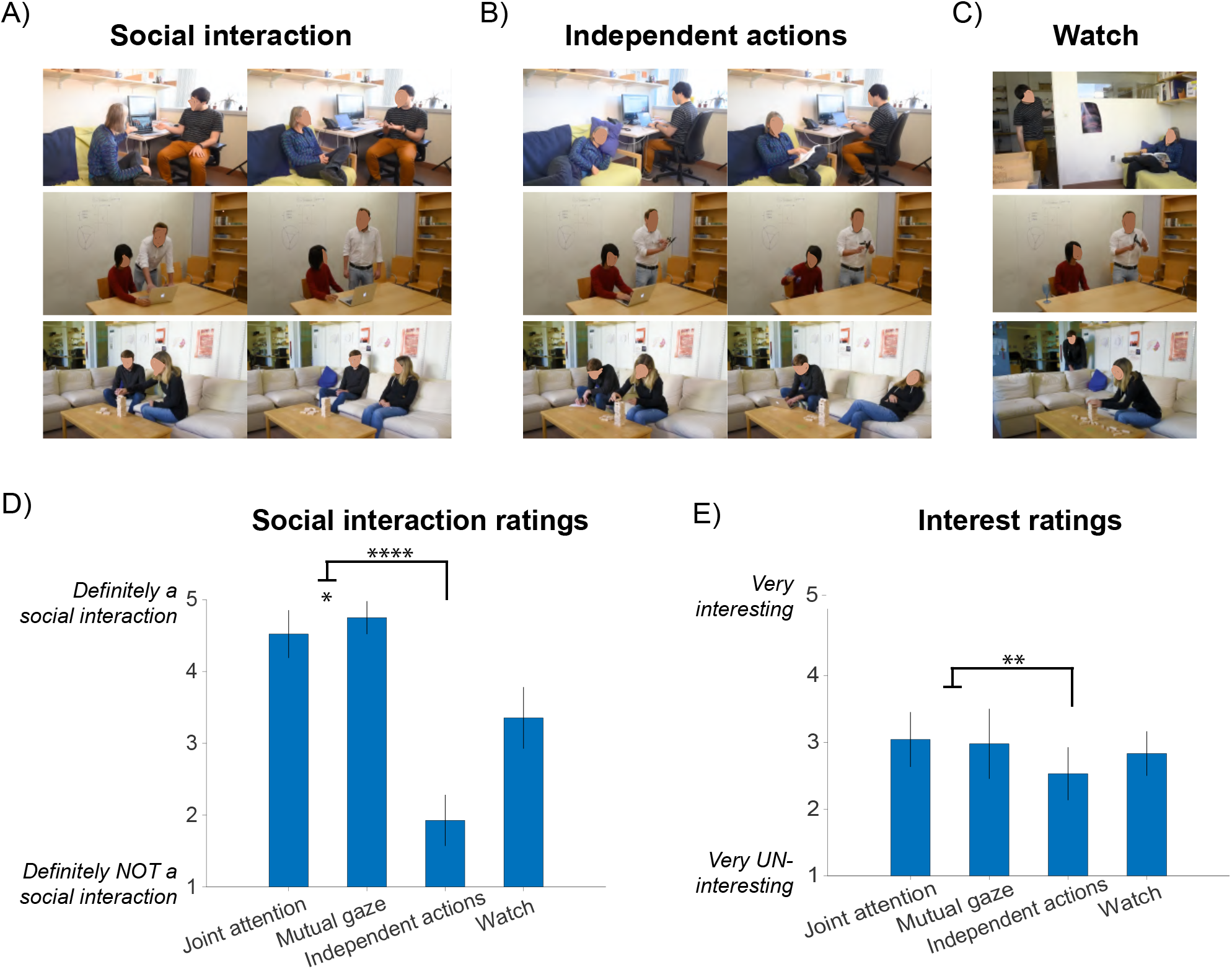
Stimulus set and behavioral responses. Subjects in the MEG viewed images depicting two actors engaged in (A) a social interaction, (B) independent actions, or (C) one actor watching the second. (Note faces have been obscured in figure to comply with biorxiv policy.) The social interaction images consisted of either a joint attention event (A, left column) or a mutual gaze event (A, right column). After the experiment, subjects rated (D) the extent to which each image depicted a social interaction (from 1 = “definitely not” to 5 = “definitely”), and (E) the visual interest of each image (from 1 = “very un-interesting” to 5 = “very interesting”). Error bars show standard deviation across subjects.

During MEG recording, each subject viewed each of the 60 images 30 times, randomized within block, while performing an orthogonal task. In particular, subjects were asked if the two actors were of the same or different gender. This task was balanced across actor pairs and scenes and the presence versus absence of social interactions. First, to replicate prior visual decoding studies and to ensure data quality, we asked whether we could decode the 60 individual images based on subjects’ MEG signals. These images included different scenes and actors, and hence differ in many visual properties. We trained a linear classifier on the MEG response at each 10 ms time bin on 80% of the trials, and tested it on the remaining 20%. We found that we could significantly decode which image subjects viewed beginning at 60 ms after image onset (**Figure 2A**). This time course of image decoding replicates several prior MEG decoding studies (Carlson et al., 2013b; Cichy et al., 2014; Isik et al., 2014), and presumably reflects primarily early visual processing. We next trained and tested a classifier at each training timepoint and each testing timepoint to generate a matrix of decoding accuracies across all train and test time points (King and Dehaene, 2014; **Figure S2A**). Decoding accuracy was highest on the diagonal, when the classifier was trained and tested at the same time point, and was only significant during a narrow time window around the diagonal. This finding suggests that the neural signals are highly dynamic, in line with previously reported results of visual decoding (Cichy et al., 2014; Isik et al., 2014; Zhang et al., 2011).

**Figure 2:**
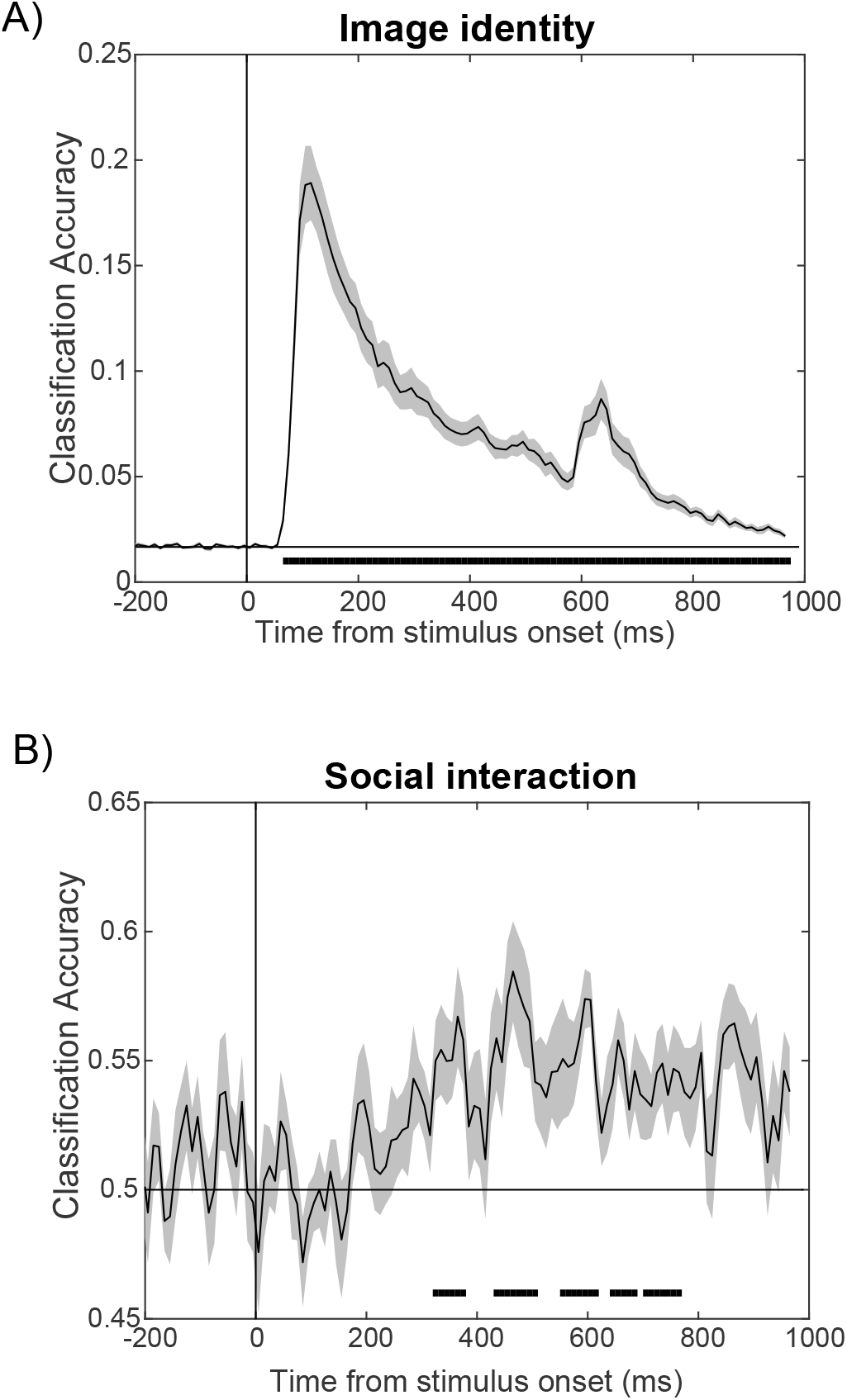
Image identity and social interaction decoding from MEG signals in Experiment 1. (A) time series of 60-way image identity decoding, with significant onset at 60 ms. (B) time series of social interaction vs. independent images decoding, with significant onset at 320 ms. (n = 16 subjects; error bars indicate SEM; vertical line indicates stimulus onset; black lines below time series indicate significant time points; two-sided permutation test; p < 0.05 cluster defining threshold; p < 0.05 cluster threshold).

We next asked the central question of this experiment: when do MEG signals encode information about whether a scene contains a social interaction (joint attention and mutual gaze conditions vs. independent action conditions)? To obtain abstract representations of social interactions, invariant to visual scene and actor information, we trained our classifier on data from subjects viewing 10 of the 12 scenarios and tested on the two held-out scenarios. This is a strong test of generalization, as the images within each scenario are much more visually similar to each other than they are to the other images in the dataset. We found that we could indeed read out the presence versus absence of a social interaction invariant to scene (**Figure 2B**). This readout occurred relatively late however, beginning at 310 ms after stimulus onset. Applying the same temporal generalization approach as before, we found again a primarily diagonal decoding pattern, indicating transient social interaction representations (**Figure S2B**). These results suggest that humans spontaneously form abstract representations of social interactions, but this occurs later than the time scale of primarily feedforward processes as in the case of invariant object recognition.

#### Social interaction decoding cannot be explained by visual interest or eye movements

We next asked if other experimental factors could account for this social interaction decoding. First, subjects rated the social interaction images as slightly more visually interesting than the noninteracting images, but this was not uniformly true across image pairs. In an exploratory analysis (not included in our pre-registration), we took the half of the image pairs with the smallest difference in interest ratings between the social and non-social images. We found that although there was no longer a significant difference in subjects behavioral interest ratings (mean rating 2.9±0.44 and 2.8±0.35, p = 0.29), we could still decode scenes with vs. without a social interaction (**Figure S3**). Overall the timecourse of decoding looked very similar to that for all images (though the onset of significant decoding did not occur until 400 ms, likely due to lower power). Thus, differences in generic attention or interest are unlikely to account for our ability to decode the presence of social interactions.

We next asked if subjects’ eye movements varied systematically across interacting and non-interacting images. To do this, we followed the same decoding procedure as with the MEG data, but instead used subjects x,y eye-position as input to our classifier. We found that we could indeed decode the presence of social interactions based on subjects’ eye position (**Figure S4A**). To address this alternative account of our findings, in a second experiment (see below) we presented the images at a smaller visual angle and for a shorter duration. While we were still able to decode scenes with vs. without a social interaction based on subjects’ MEG data, we could no longer do so based on eye position (**Figure S4B**).

#### Feedforward computational model cannot recognize social interactions across scenes

To more explicitly test whether feedforward computations can distinguish between scenes with vs. without a social interaction, we asked if a purely feedforward deep convolutional neural network model (VGG-16, trained on Imagenet (Simonyan and Zisserman, 2014)) could distinguish between these two conditions. While this model was trained on an object recognition task, it has seen a wide range of natural images, and similar networks have been shown to generalize across different visual pattern recognition tasks (Blauch et al., 2017; Zhou et al., 2014). We used the response of each unit in each layer of the CNN as features to our linear classifier. Just as with the MEG data, we trained our classifier on 10 of the 12 social and non-social scenes, and tested on the two held out scenes. We could not detect the presence vs. absence of a social interaction, in a manner that generalized across scenes, with the output of any layer of the CNN (**Figure 3**, red bars). In the final pooling layer of the model, we could detect social interactions in a manner that did not generalize across scenes (**Figure 3**, blue bars). This was analogous to our MEG data where we detected the presence of a social interaction earlier, within 200 ms, in random 80-20 splits of our data (requiring no generalization across scenes, **Figure S1**). Taken together, these results suggest that standard feedforward computations are not sufficient to detect abstract social interactions in natural scenes.

**Figure 3:**
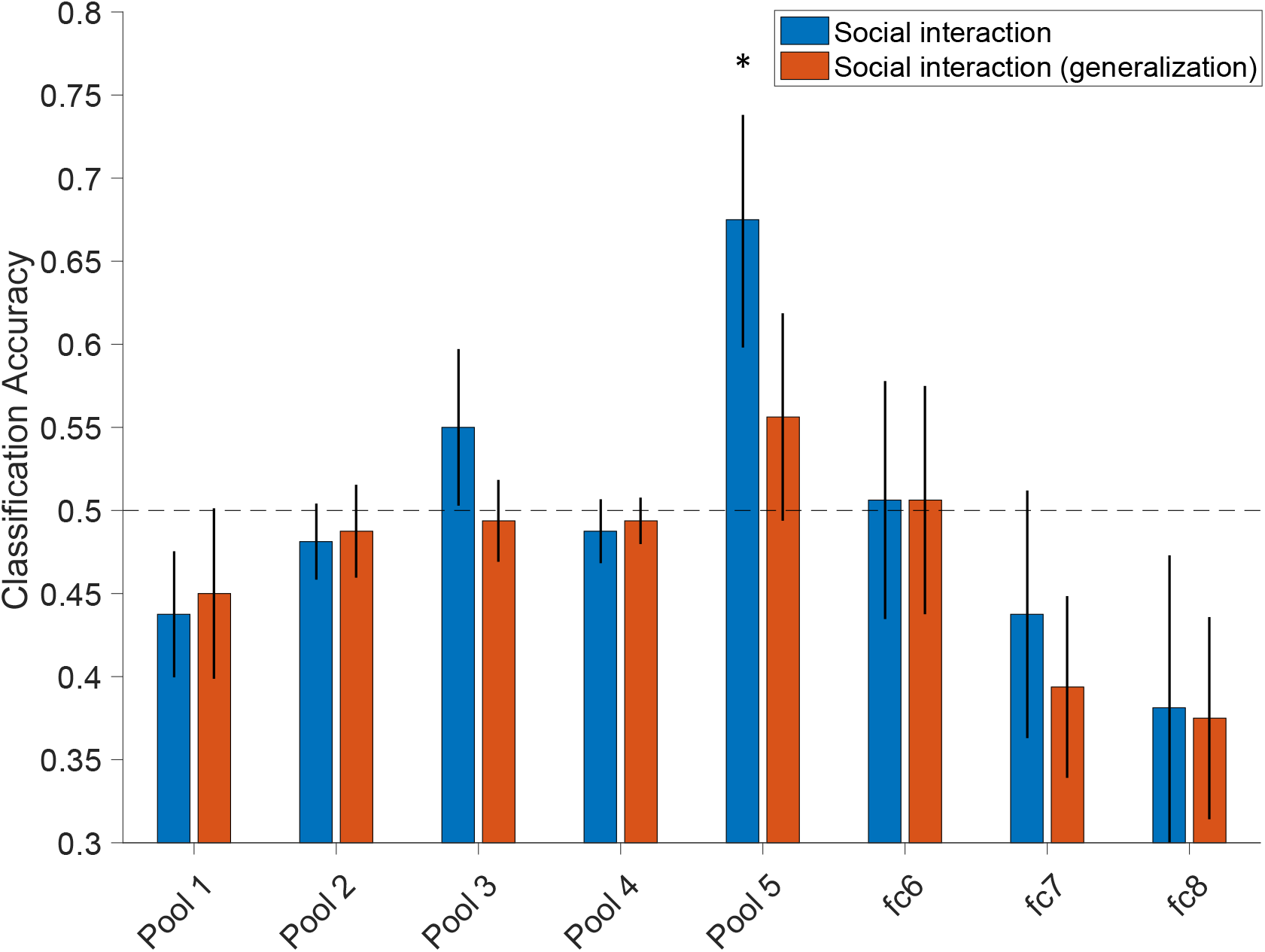
Social interaction decoding from different layers of a convolutional neural network model. Bar plot shows the classification accuracy of each layer for the output of each layer in a social interaction detection task. Blue bars indicate decoding without generalization, and red bars decoding with generalization across different scene images. Error bars indicate SD across multiple resample runs. Asterisks indicates p<0.05 significant decoding based on permutation test.

#### Distinct representation of two-way social interactions

We next asked what information is driving our ability to decode social interactions. Is it sufficient for one agent to be aware of the other (“perceptual access”), or is an actual two-way social interaction necessary? To answer this question we asked if we could decode one-way “watch” images (in which one agent sees the other but not vice versa) from two-way “mutual gaze” images. We found that we could decode watch versus mutual gaze at similar latency to social interaction read out (**Figure 4a**, onset 330 ms). Note that this decoding is based on half as much data as the social interaction detection analysis above, so the read out is necessarily noisier. Interestingly, we could not decode watch images from independent action images (**Figure 4b**). These findings suggest that it is the presence versus absence of a two-way social interaction that is a distinct and spontaneously represented property of the image, not the mere presence of one-way perceptual access from one agent to the other.

**Figure 4:**
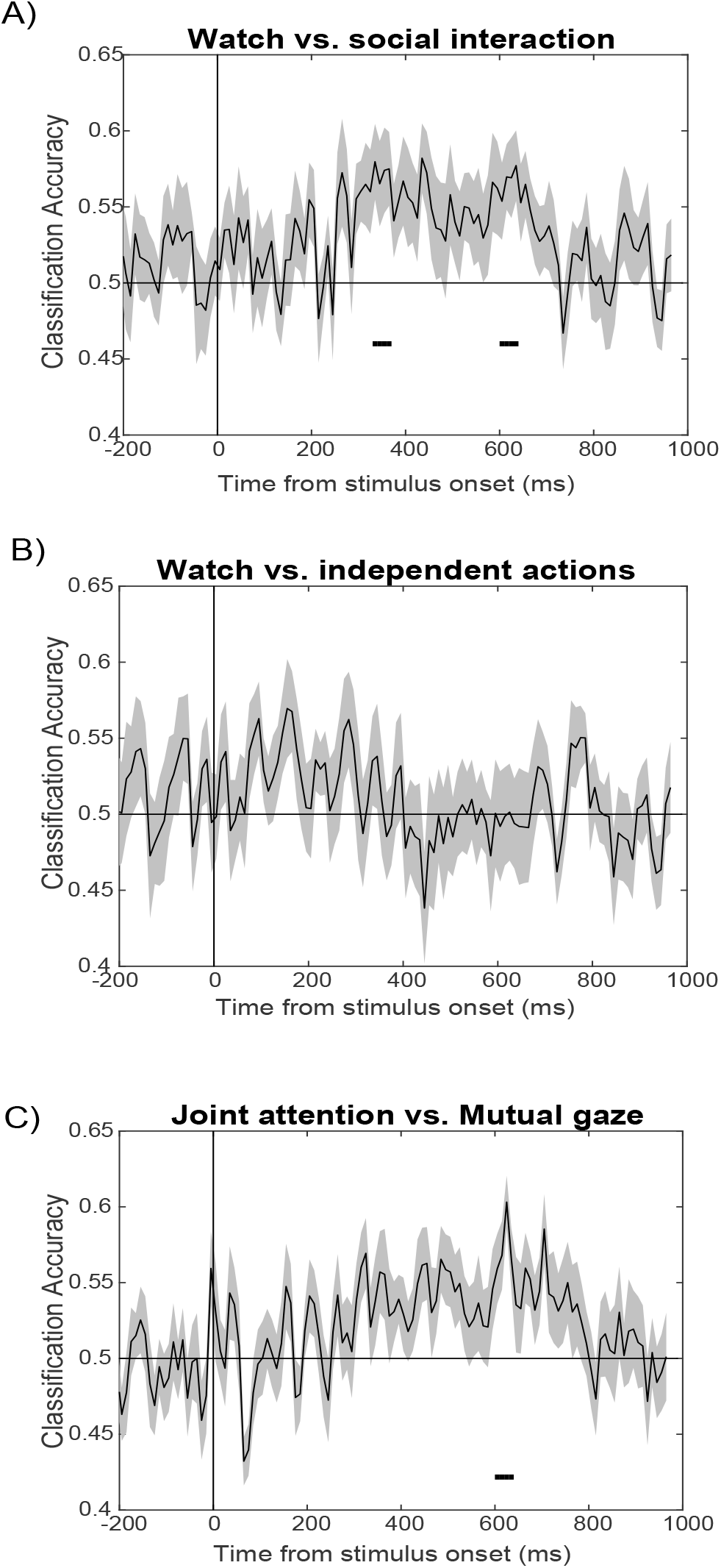
Decoding different types of social interactions from MEG signals in Experiment 1: Time series of (A) watch vs. social intearction (onset=330 ms), (B) watch vs. independent actions (not signifcant), (C) mutual gaze vs. joint attention (onset= 600 ms). Error bars and statistical tests are the same as in **Figure 2**.

#### Late readout of Type of Third Party Social Interaction

Finally, beyond simply detecting the presence of a social interaction, we asked whether the MEG signal contained information about the different type of social interactions in our dataset: Joint attention vs. Mutual Gaze. Mutual gaze is perhaps the most perceptually obvious form of social interaction between two agents. But joint attention is also a fundamental form of social interaction that arises early in infancy (Scaife and Bruner, 1975) and may be critical in language learning (Tomasello and Farrar, 1986). Do perceivers spontaneously distinguish between these two forms of social interaction, and if so, when? We found that we could distinguish joint attention vs. mutual gaze in a manner that generalized across scenes, but only quite late, at 600 ms after image onset (**Figure 4c**). This analysis makes use of only half the data as the analysis of social interaction detection and thus could fail to detect earlier discriminative information (but see Experiment 2/**Figure 6** for a replication of these results).

In sum, the results of Experiment 1 suggest that both the presence and type of social interactions are spontaneously represented in the brain, but this information comes online very late, well beyond the timescale of primarily feedforward processes.

### Experiment 2

#### Social interaction perception is slow even during an explicit task

In a second experiment we asked if there were any conditions under which social interaction information could be extracted more quickly. In particular, can these neural computations occur faster if the subject is explicitly asked to behaviorally extract that information? This would be consistent with prior studies showing rapid visual readout that also used an explicit task (Thorpe et al., 1996). To test this hypothesis, we ran a second experiment with 16 additional naïve subjects who saw identical images in the MEG, but now instead of performing the same versus different gender task they performed an explicit social interaction task (i.e., does this image contain a social interaction?). The button order was flipped halfway through the study so that explicit motor responses could not account for our decoding results (see Methods). In this experiment, we removed the “Watch” stimuli to for this experiment because they are ambiguous in terms of whether they should or should not count as a social interaction, and they are not directly relevant to our central question. Subjects’ mean reaction time on the social interaction task was 1.0±0.25 s (mean±SD across subjects). As expected, we found no difference in the onset latency of image identity decoding (**Figure 5A**).

**Figure 5:**
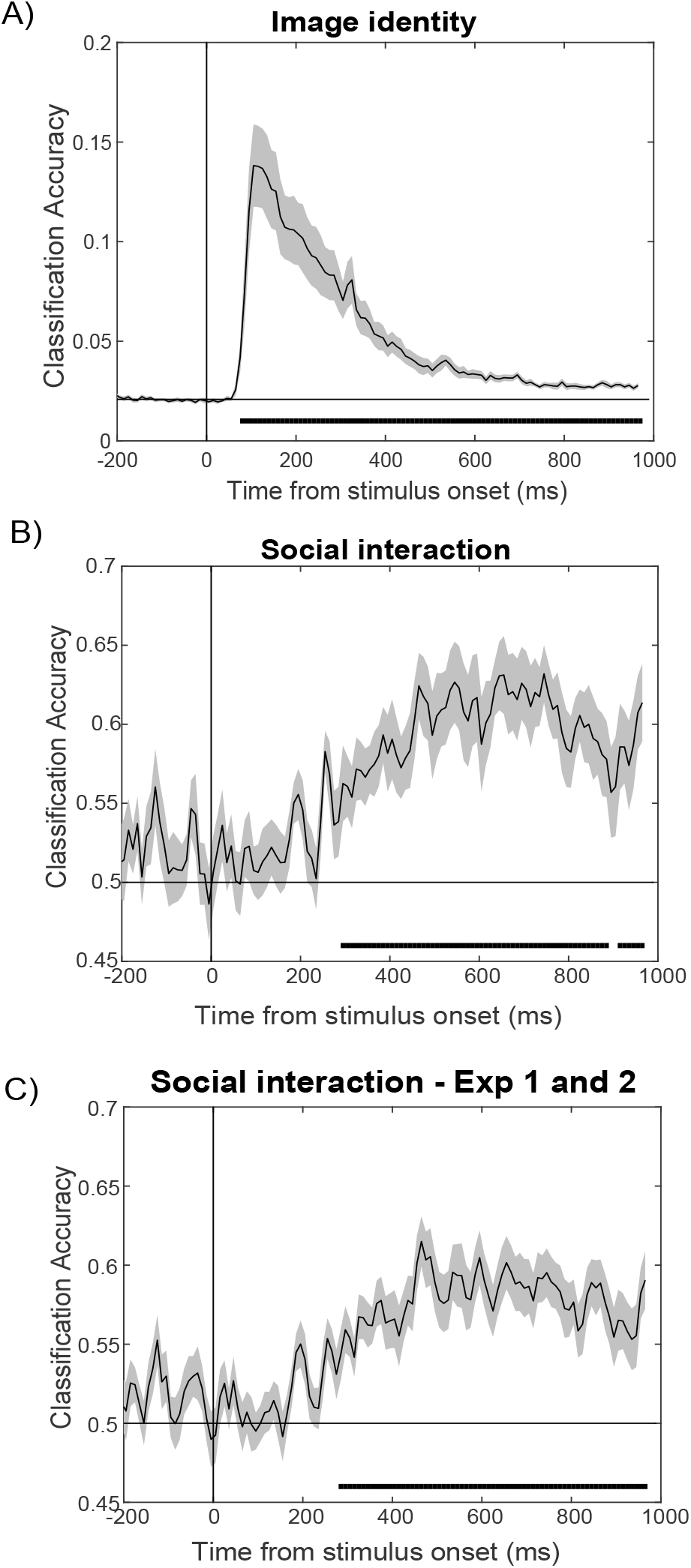
Image identity and social interaction decoding from MEG signals in Experiment 2: Time series of (A) 48-way image identity (onset = 70 ms), (B) social interaction vs. independent images generalizaing across scenes (onset = 300 ms), (C) combined Exp 1 and 2 (onset = 290 ms). Error bars and statistical tests are the same as in **Figure 2**.

Importantly, we found that even when subjects performed an explicit social interaction detection task, information about the presence vs. absence of a social interaction could again only be discriminated in MEG response at relatively late latencies after stimulus onset (290 ms, **Figure 5B**). Could earlier social interaction signals exist that we did not have sufficient power to see in either experiment? To answer this question, in an exploratory analysis, we combined the data from Experiments 1 and 2 and re-ran our social interaction decoding analysis. Even with 32 subjects, the onset latency of social interaction detection did not change (**Figure 5C**).

Next, we again asked if and when we could decode the type of social interaction (mutual gaze vs. joint attention) in Experiment 2, as we could in Experiment 1. While the latency moved slightly earlier, it was still quite late, with an onset of 530 ms after stimulus onset (**Figure 6A**). In another exploratory analysis, we combined data across Experiments 1 and 2 and found that in all 32 subjects onset latency was 490 ms (**Figure 6B**).The results of Experiment 2 serve as an internal replication, and confirm that even in the presence of task demands, social interaction perception is computed well beyond the timescale of visual pattern recognition.

**Figure 6:**
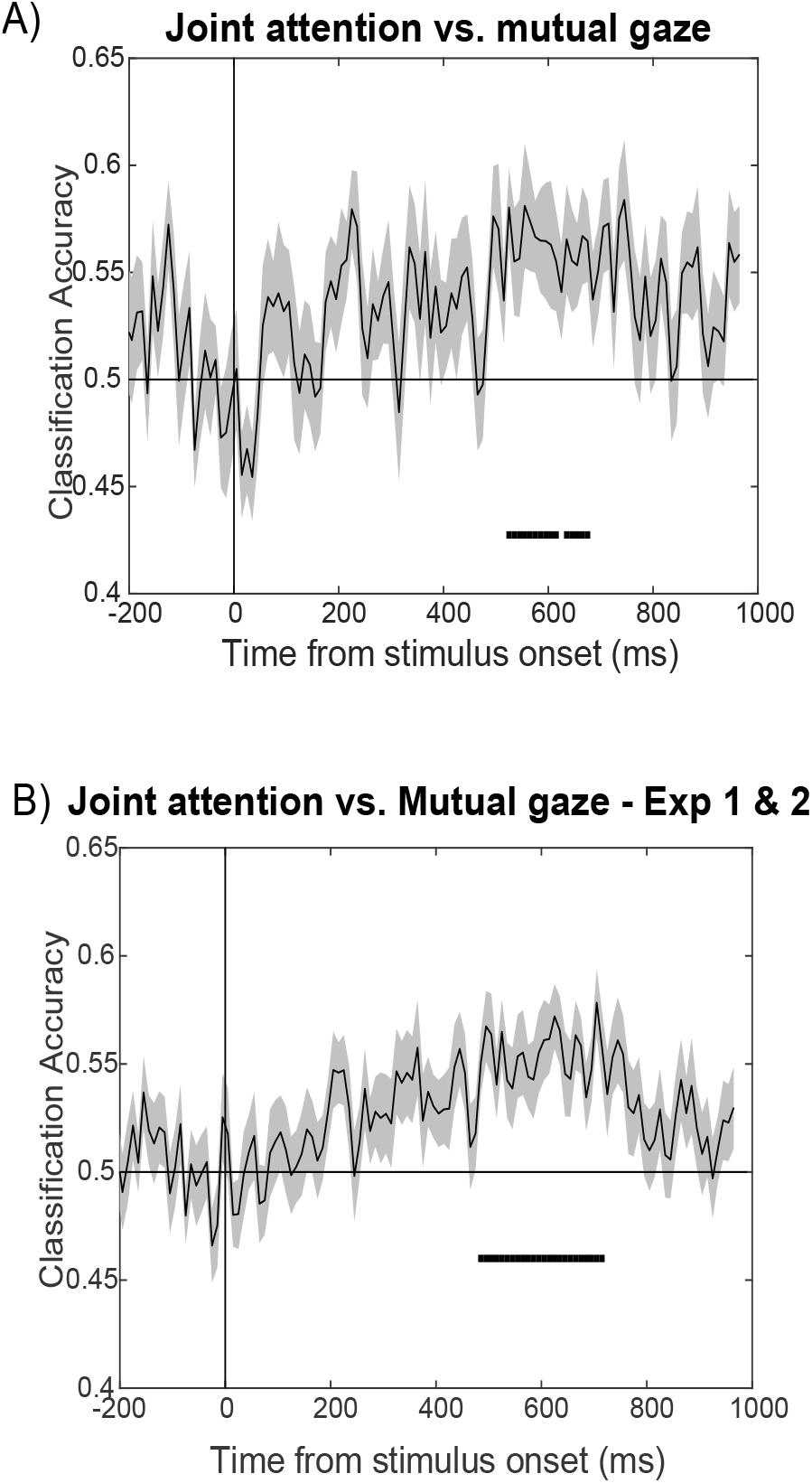
Decoding different types of social interactions from MEG data in Experiment 2. Time series of mutual gaze vs. joint attention for (A) Experiment 2 (onset 530 ms), and (B) combined Exp 1 and 2 (onset 490 ms). Error bars and statistical tests are the same as in **Figure 2**.

#### MEG-Behavioral correlation

As with all decoding studies, it is important to investigate whether the neural information we extract is associated with perceptual judgements, or whether it is merely epiphenomenal (Grootswagers et al., 2018; Williams et al., 2007). One way to address this concern is to test whether readout performance is tied to subjects behavioral judgments and reaction times. Because we had these measures in our second experiment, we correlated each subjects MEG and behavioral data using representational similarity analysis (RSA). Specifically, we computed the time-resolved dissimilarity matrix for our MEG data (48×48 pairwise image decoding accuracy) and a behavioral dissimilarity matrix (48×48 behavioral dissimilarity matrix), and correlated these measures within subject. We found that there was indeed a significant correlation between subjects’ behavioral accuracy and MEG data beginning 340 ms after stimulus onset (**Figure 7**). These results suggest that the MEG signals we detected are behaviorally relevant to social interaction perception.

**Figure 7:**
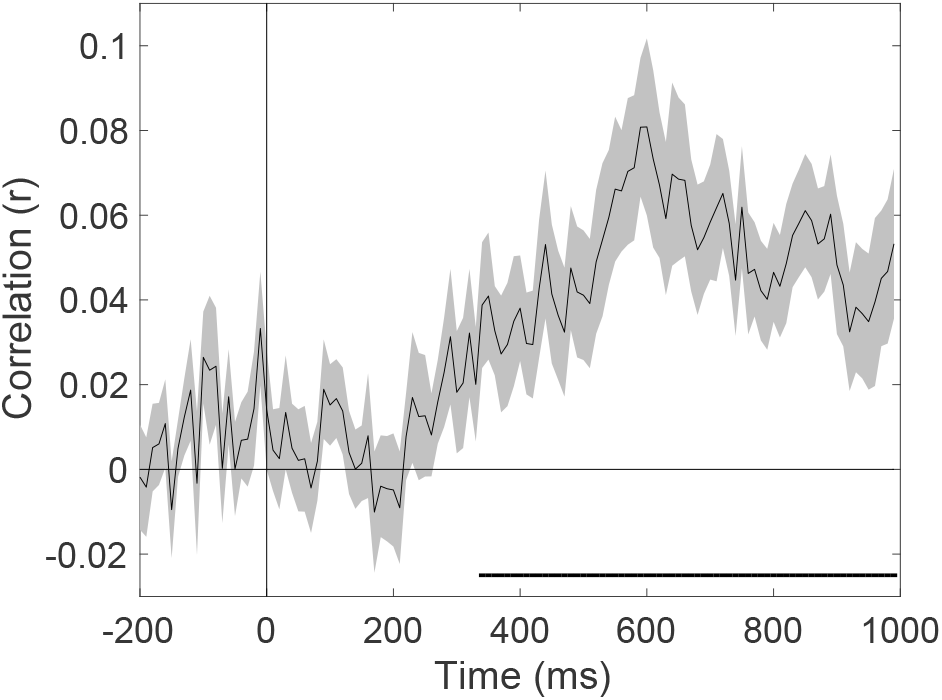
Social interaction information encoded in MEG signals was correlated with behavior. Time series shows the correlation between the MEG dissimilarity matrix at each time point and reaction time dissimilarity matrix. Error bars and statistical tests are the same as if **Figure 2**.

## Discussion

In this work, we identified neural signals that contain information about the presence and type of third-party social interactions in a visual scene. These neural representations generalized across low-level visual features, and arose spontaneously, even when participants performed an orthogonal task. Crucially though, they arose relatively late compared to previously reported latencies for other types of visual pattern classification: 300 ms after stimulus onset for detection and 600 ms for categorization. These late latencies were found even when subjects performed an explicit social interaction detection task. Importantly, this neural readout was correlated with behavior. In line with this late onset time, we found that a standard feedforward neural network could not distinguish between scenes with versus without a social interaction. Taken together, these results suggest that social interaction perception is not based on purely feed-forward processing, but instead relies on slower, presumably recurrent computations.

The latencies of social interaction detection and categorization (300 and 600 ms, respectively) are substantially later than those previously reported for different types of pattern recognition problems, such as invariant object recognition (Carlson et al., 2013a; Isik et al., 2014; Yamins et al., 2014). Prior work with ERPs (Thorpe et al., 1996) and physiology (Yamins et al., 2014) indicates that object recognition is quite fast, occurring between 100-200 ms after stimulus onset, even when the objects appear in complex backgrounds, like the natural images used in our study. Natural scene information has been shown to arise on a similarly fast time scale to object recognition (Cichy et al., 2016a; Greene and Hansen, 2018).

In contrast to object and scene perception, there has been relatively little M/EEG decoding work on aspects of social perception. The N170 response (Bentin et al., 1996) is a strongly face-selective univariate response arising around 170 ms after image onset. However, recent decoding studies have shown that many aspects of face information are represented earlier than 170 ms. For example, age, gender and identity are all decodable around 100 ms (Dobs et al., 2018). Even emotion properties like expression (100 ms (Dima et al., 2018)) and valence and arousal (150 ms (Grootswagers et al., 2017)) have been shown to come online quickly. Beyond face properties, the emotional valence and self-relevance of communicative gestures can be decoded within 100 ms (Redcay and Carlson, 2015), and individual agents’ actions as early as 200 ms (Isik et al., 2018). Interestingly, like object and scene perception, these social dimensions all fall within the rough timescale of feedforward pattern recognition. The present study suggests that the detection of third-party social interactions occurs substantially later, and thus may be based on fundamentally different computations from these other visual and social recognition processes. A critical difference may be that, unlike face, emotion, action, and gesture recognition of individuals, social interaction recognition involves taking into account relational information between multiple agents.

In general, feedforward models perform poorly on tasks that involve incorporating relational information (Yuille and Kersten, 2006). It is important to note, however, that while we showed that a generic deep neural network trained on object classification could not detect scenes with vs. without social interactions, we do not know how a similar neural network trained on a social interaction task would perform. Unfortunately, this question is difficult to answer due to the lack of large-scale labelled image datasets in this domain. However, similar feedforward networks have been shown to generalize across different types of pattern recognition problems. For example, networks trained on either scene or object recognition perform well above chance on the opposite problem (Zhou et al., 2014). Similarly, a network trained on face recognition has been shown to generalize somewhat to both object and scene recognition problems (Blauch et al., 2017). In contrast this network, trained on an object recognition task, did not perform above chance in our social interaction task (**Figure 3**). While it is difficult to rule out earlier feedforward signals or computations that we could not detect with MEG or an appropriately trained model, convergent evidence from these two methods suggests that social interaction detection cannot be solved with fast, feedforward computations alone. It remains an interesting open question whether a model with recurrence, which largely has same structure as the deep neural network tested here, would perform well on the task, or if a fundamentally different class of generative models is required to solve this problem.

This study takes just a first step toward understanding the time course and computations underlying multi-agent social interaction perception, but many questions remain. Beyond detecting simple, dyadic social interactions and distinguishing between different gaze events, what other social interaction information is spontaneously extracted by the brain? How do our brains code complex real-world complex social events such as a party or sporting event? And how fine-grained are the automatically-extracted representations of social interactions (positive vs. negative, different action categories, etc.)? And perhaps most obviously, where in the brain do these social interaction representations originate? Our prior fMRI results indicate that a region in the right posterior superior temporal sulcus is selectively engaged in the perception of social interactions (Isik et al., 2017). If this region does indeed underlie the decoding information we report here with MEG, does it receive input from purely from visual regions, or from higher level regions that code for information about individual social agents (Grossman et al., 2000; Puce et al., 1998; Saxe and Kanwisher, 2003) or the physical world around them (Fischer et al., 2016)? While future work combining fMRI and MEG will be needed to answer these questions, this work provides important initial constraints on the neural computations underlying multi-agent social interactions.

## Methods

### Social interaction dataset

We created an image dataset depicting pairs of people interacting with each other or independently in different ways. There were five different conditions, shot across 12 scenes with 12 different actor pairs (60 images total, see **Figure 1A** for example images). The five conditions differed in the way each pair of people were or were not interacting with each other. The different conditions include:

i. Mutual gaze – pair of actors is looking at each other. Social interaction.
ii. Joint attention – pair of actors looking at the same object. Social interaction.
iii. Independent actions 1 – two actors are engaged in separate independent actions. No social interaction.
iv. Independent actions 2 – two actors are engaged in separate independent actions (different actions from above). No social interaction.
v. Watch – one actor watches the other actor who is looking away. One-way interaction.

### Subjects

32 naïve subjects (16 for Experiment 1, and 16 different subjects for Experiment 2) between 18-45 years old with normal or corrected to normal vision participated in these experiments. Our experimental protocol was approved by the MIT Committee for the Use of Humans as Experimental Subjects. Three additional subjects were excluded from Experiment 1 based on a pre-defined behavioral exclusion criteria (<80% accuracy on behavioral task).

### Experimental procedure

#### Experiment 1

Subjects viewed the 60 images presented 30 times each in the MEG. The order of the 60 images was randomized within each block. Subjects were instructed to fixate centrally and judge if the two people in each image were the same or different genders. The images were presented at 9×5 degrees of visual angle for 500 ms each with a central fixation cross. Each image was immediately followed by task instructions, and subjects responded yes or no. After each question a 200ms fixation cross would appear before the next image. Task responses were self-paced, and the button order flipped halfway through the experiment to avoid motor confounds.

#### Experiment 2

In Experiment 2 the procedure was exactly the same as Experiment 1 except 1) subjects viewed only 48 of the 60 images (excluding “watch” condition), 2) to mitigate the effect of eye movements images were presented smaller (5 x 2.8 degrees of visual angle) and for a shorter duration of 200ms, and 3) subjects performed an explicit social interaction task (“Are these two people engaged in a social interaction?”).

### Eye tracking

We tracked the subjects’ left and right eye positions using an Eyelink 1000 eye tracker with a 9-point calibration. We were not able to achieve an accurate calibration for 4 subjects in Experiment 1 and 3 subjects in Experiment 2, so these subjects’ eye position data were excluded from the eye tracking analysis. To test whether there is stimulus-selective information was present in our eye tracking data, we performed the below decoding procedures using the X,Y output of each eye as classifier features (see Decoding Methods for more details).

### MEG acquisition and pre-processing

The MEG data were collected using an Elekta Neuromag Triux scanner with 306 sensors, 102 magnetometers and 204 planar gradiometers, with an online bandpass filter between 0.01 and 330 Hz. Subjects head position was continuously monitored throughout the experiment using five head position index (HPI) coils. First the signals were filtered using temporal Signal Space Separation and motion corrected (based on the position of the HPI coils) with Elekta Neuromag software. Next, Signal Space Projection (Tesche et al., 1995) was applied to correct for movement and sensor contamination. The MEG data were divided into epochs from −200 to 1000 ms, relative to video onset, with the mean baseline activity removed from each epoch. The signals were band-pass filtered from 0.1 to 100 Hz to remove external and irrelevant biological noise (Acunzo et al., 2012; Rousselet, 2012). The above preprocessing steps were all implemented using the Brainstorm software (Tadel et al., 2011).

### MEG decoding

We analyzed the MEG data using the neural decoding toolbox for Matlab (Meyers, 2013). We averaged the data in each sensor into 10ms non-overlapping bins, and trained and tested a new linear correlation coefficient classifier at each time point. We used 5-fold cross validation (CV) splits (training on 80% of the data and testing on the held out 20%). We performed feature selection using an ANOVA on only the training data (to avoid double dipping/circularity) and selected the 25 sensors whose activity most significantly co-varied with the training labels. These selected sensors were fixed for testing. We repeated the entire decoding procedure at each time point 20 times and report the mean accuracy for each condition. See (Isik et al., 2018, 2014) for a more detailed description of the decoding methods.

The variables we decoded decode were:

1. 60-way image identity. Each image was repeated 30 times, and we divide the data into five cross validation splits with 6 trials per CV split. To increase signal to noise, we averaged the data from all 6 trials together.
2. Social interaction (mutual gaze and joint attention) vs. independent action images (2 non-interacting conditions per scenario).
3. Joint attention vs. mutual gaze.
4. Social interaction (mutual gaze) vs. watch.
5. Non-interacting images vs. watch.

For tests 2-5, we ran the decoding in a manner that generalized across scenario. In particular, we trained our classifier on 10 scenarios and tested on the remaining two, held-out scenarios. For the generalization decoding, we averaged 30 trials together (note that in our pre-registration we stated we would average 24 trials together. This was an error as 24 is not divisible by the number of trials included in conditions 3-5 so would require us to exclude data from the decoding).

### Statistical inference

We assessed decoding significance using non-parametric statistical tests that do not make assumptions about the underlying distribution of the data (Pantazis et al., 2005). Specifically, we performed a sign permutation test that centers each subjects’ MEG data around chance and randomly multiplies it by +1 or −1. We repeated this procedure 1000 times to generate a null distribution. To correct for multiple comparisons, we used cluster correction in time with a cluster defining threshold of p<0.05 and a corrected significance level of p<0.05 (Cichy et al., 2016b; Mohsenzadeh et al., 2018).

### CNN model

To further test if social interaction detection can be performed using feedforward computations, we ran our stimuli through a pre-trained feedforward deep neural network: VGG-16 trained on Imagenet (Simonyan and Zisserman, 2014). We asked if the output of each of the models’ five pooling layers could distinguish between images with vs. without a social interaction. First, to reduce the dimensionality of each layers’ output we performed PCA and selected the top 50 components from each layer (note our pilot data showed very similar results with 40-59 components). Within each layer, we then took the response to each image and, as with our MEG data, trained a linear classifier to distinguish between scenes with vs. without a social interaction on data from 10 of 12 scenes. We tested the linear classifier on data from the two held-out scenes. We repeated this procedure 20 times, holding out two random scenes each time.

### Representational similarity analysis

We compared our MEG data to our behavioral data in Experiment 2 using representational similarity analysis (RSA; Kriegeskorte et al., 2008). To produce the MEG dissimilarity matrix, we followed a similar procedure to (Cichy et al., 2014). We first performed PCA on the MEG responses to the 60 images in our data to reduce the dimensionality of the data the number of components that explains 99.99% of original variance in data. We next calculated the dissimilarity between each pair of images based on their pairwise classification accuracy computed over those PCs. We repeated this at each time point to get a new dissimilarity matrix.

To produce the behavioral dissimilarity matrix we used a behavioral metric that took into account the subjects’ reaction time scaled by their response. In particular, we calculated the metric as: Response*(1-RT/max(RT)), where RT is reaction time, and the response is +1 for social interactions and −1 for non-social interactions. These values ranged from +1 for the fastest social interaction responses to-1 for the fastest non-social interaction responses. For each subject, we calculated the average pairwise difference between each image pair to construct our behavioral dissimilarity matrix.

## Supplemental Figures

**Figure S1:**
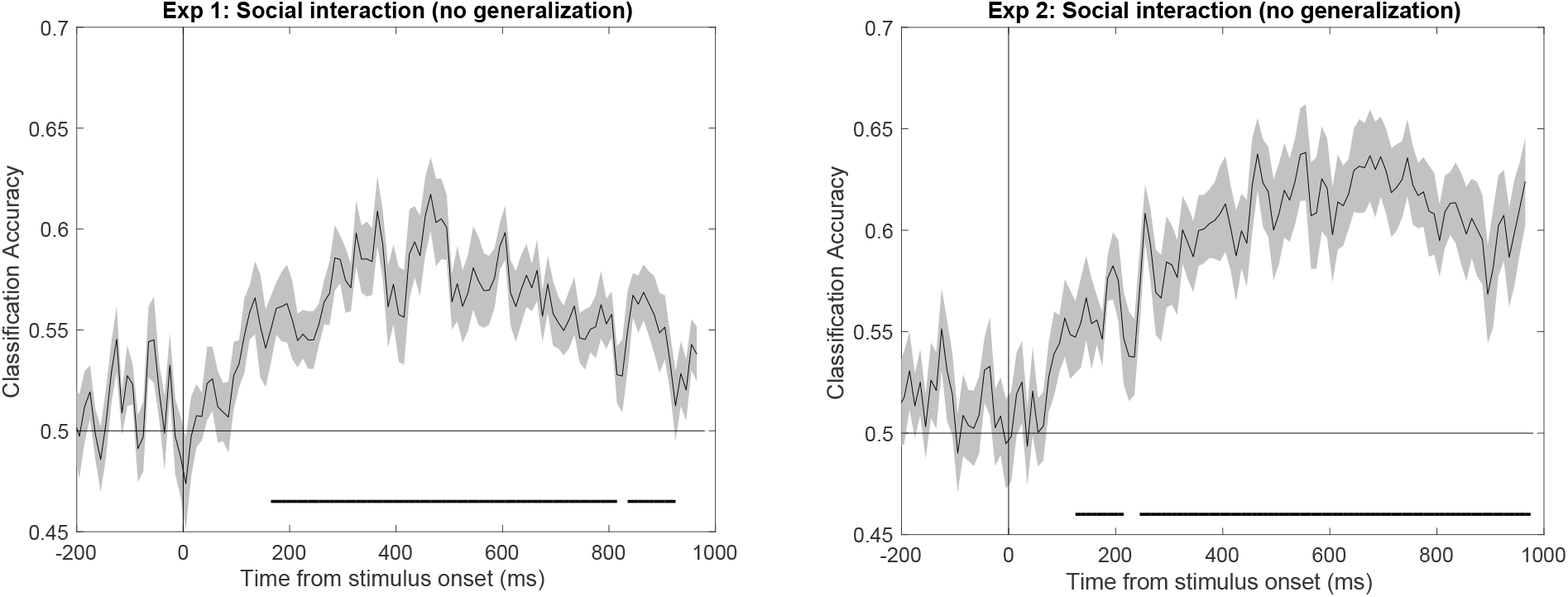
Social interaction decoding without generalization. Decoding time series of scenes with vs. without a social interaction from random 80-20 splits across all trials (possibly including different trials of the same images or scenes across spilts) and thus not explicitly testing generalizing across scenes for (A) Experiment 1 and (B) Experiment 2. error bars indicate SEM; vertical line indicates stimulus onset; black lines below time series indicate significant time points; two-sided permutation test; p < 0.05 cluster defining threshold; p < 0.05 cluster threshold).

**Figure S2:**
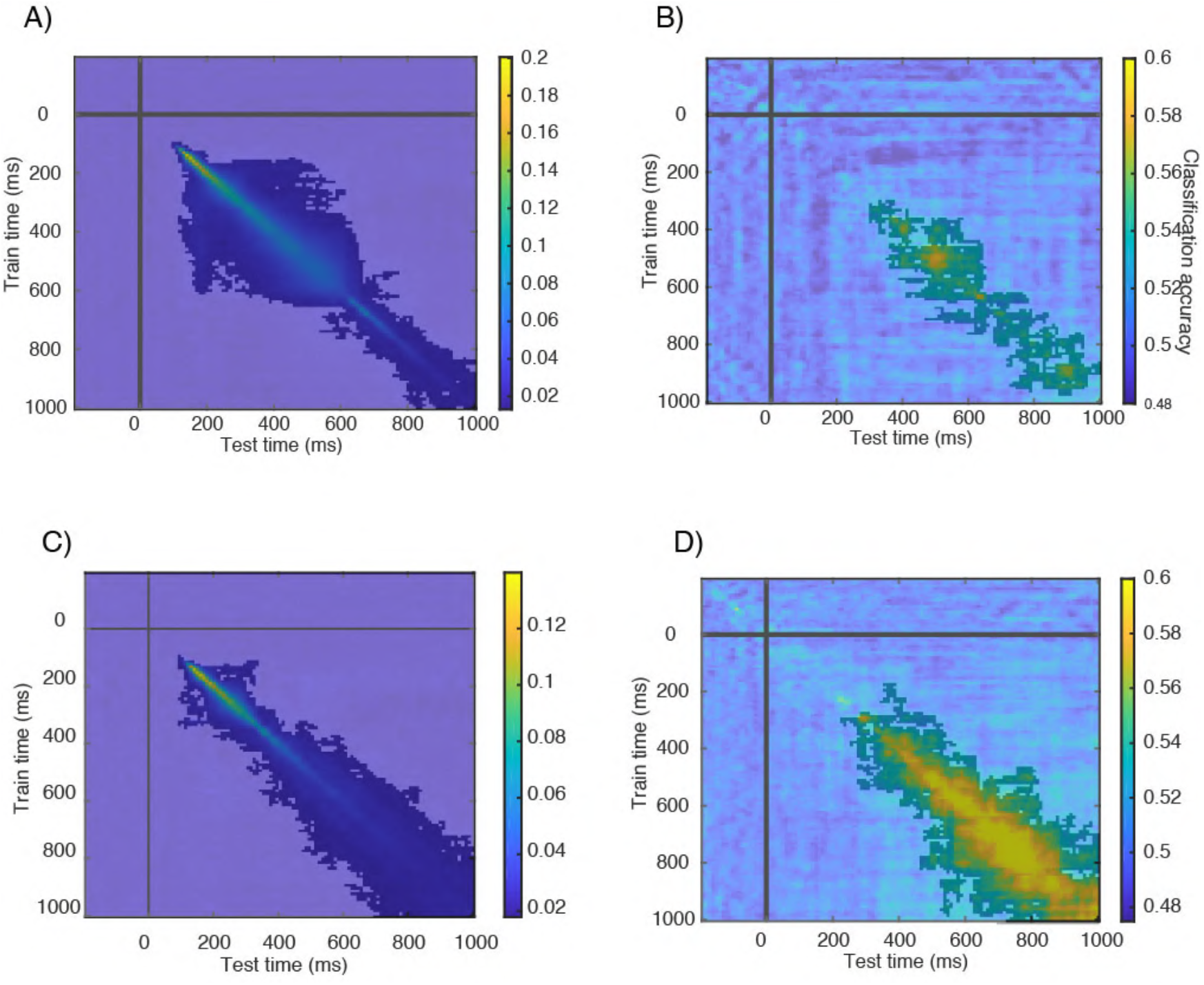
Temporal generalization decoding. Classification accuracy for training and testing a classifier at all pairs of time points (diagonal corresponds to line plots shown in **Figures 2** and **5**) for (A) Experiment 1 image identity decoding, (B) Experiment 1 social interaction decoding, (C) Experiment 2 image identity decoding, and (D) Experiment 2 social interaction decoding. Significance testing is the same as in **Figure S1**.

**Figure S3:**
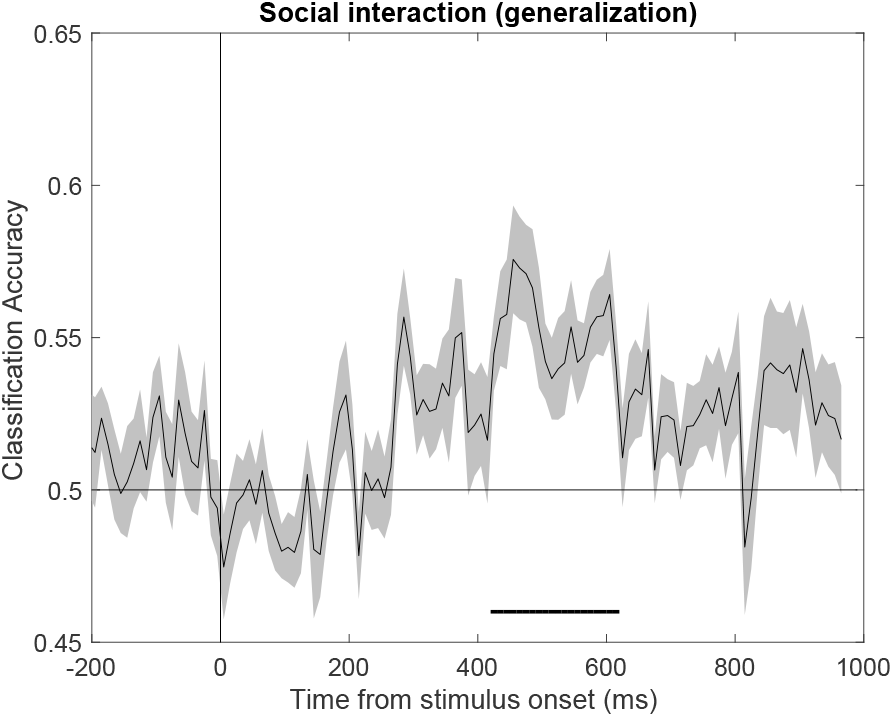
**Social interaction decoding for images matched on “interest” ratings** in post-MEG experiment (see **Figure 1F** for ratings). Error bars and significance testing is the same as in **Figure S1**.

**Figure S4:**
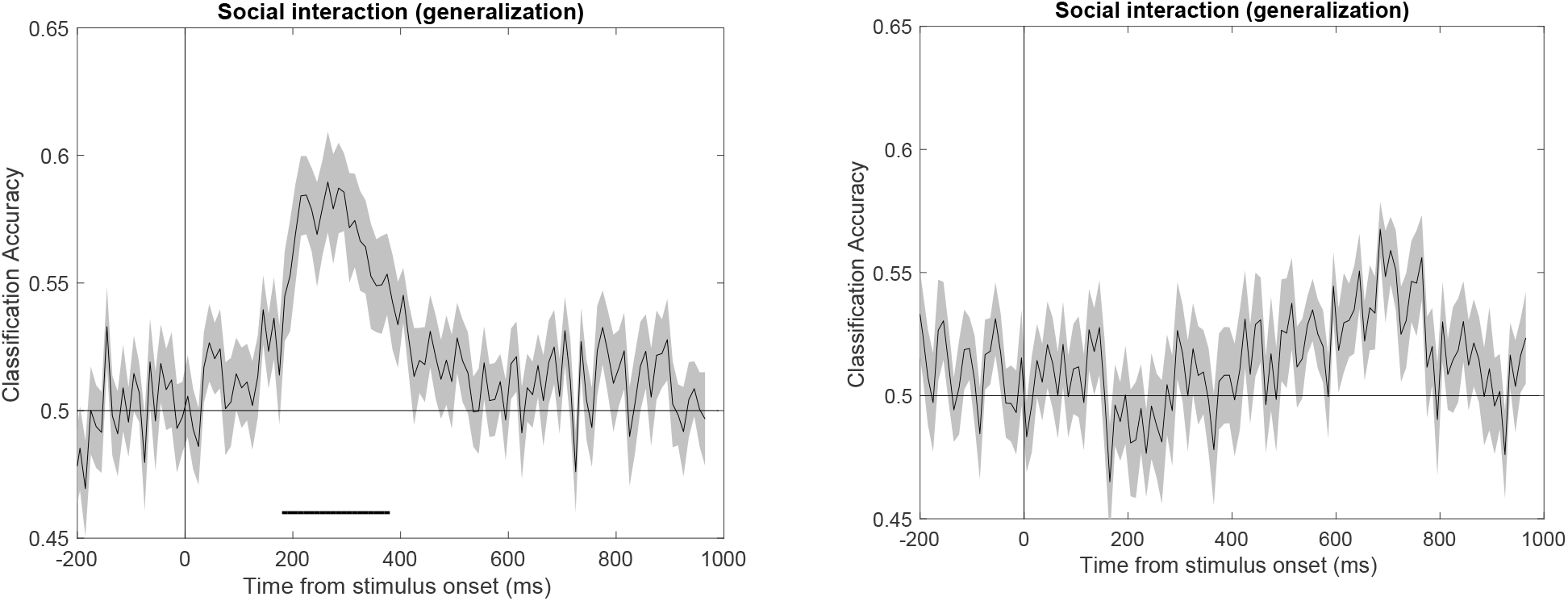
Social interaction decoding from eye tracking signals. Time series of social interaction vs. independent images decoding in (A) Experiment 1, and (B) Experiment 2. Error bars and significance testing are same as in **Figure S1**.

